# Implementation of pupylation-based proximity labelling in plant biology reveals regulatory factors in cellulose biosynthesis

**DOI:** 10.1101/2024.01.09.574788

**Authors:** Shuai Zheng, Ouda Khammy, Lise Charlotte Marie Noack, Andreas De Meyer, Ghazanfar Abbas Khan, Nancy De Winne, Dominique Eeckhout, Daniel van Damme, Staffan Persson

**Affiliations:** Copenhagen Plant Science Center, Department of Plant & Environmental Sciences, University of Copenhagen, Frederiksberg C, 1871, Denmark; Department of Plant Biotechnology and Bioinformatics, Ghent University, Technologiepark 71, 9052 Ghent, Belgium; VIB Center for Plant Systems Biology, Technologiepark 71, 9052 Ghent, Belgium; Department of Animal, Plant and Soil Sciences, School of Agriculture, Biomedicine and Environment, La Trobe University, Bundoora, VIC 3086, Australia; Joint International Research Laboratory of Metabolic and Developmental Sciences, State Key Laboratory of Hybrid Rice, School of Life Sciences and Biotechnology, Shanghai Jiao Tong University, Shanghai 20040, China

## Abstract

Knowledge about how and where proteins interact provides a pillar for cell biology. Biotin-related proximity labelling approaches are efficiently monitoring protein interactions but may have labelling-related drawbacks. Here, we introduce pupylation-based proximity labelling (PUP-IT) as a tool for protein interaction detection in plant biology. We show that PUP-IT readily confirmed and extended protein interactions for several known protein complexes, across different types of plant systems. To further demonstrate the power of PUP-IT, we used the system to identify protein interactions of the protein complex that underpin cellulose synthesis in plants. Apart from known complex components, we identified the ARF-GEF BEN1 (BFA-VISUALIZED ENDOCYTIC TRAFFICKING DEFECTIVE1). We show that BEN1 contributes to cellulose synthesis by regulating the trafficking of the cellulose synthesis protein complex between the trans-Golgi network and the plasma membrane. Our results, therefore, introduce PUP-IT as a new and powerful proximity labelling system to identify protein interactions in plant cells.

---

Protein-protein interactions (PPIs) are at the core of all cellular processes and have the capacity to infer functions of unknown proteins. PPI analyses are done through a plethora of methods, much of which are performed in heterologous hosts or *in vitro*^1^. By contrast, the assessment of PPIs in native hosts is largely restricted to different affinity-based purification schemes or what is referred to as proximity labelling. Proximity labelling may be done by fusing a protein of interest, referred to as bait, with an enzyme that can label proteins in the vicinity of the bait. The most frequently used proximity labelling entails various versions of biotin ligases or peroxidases ^2,3^. Some of the drawbacks of these systems include endogenous enzyme activities that may confound the results and substrate availability^2,3^. However, recent developments have produced completely genetically encoded proximity labelling systems, such as pupylation (PUP-IT; Fig. 1a)^4^. PUP-IT is proposed as a promising proximity labelling system in plant cells but has so far only been used in transient protoplast assays^5^.

**Figure 1:**
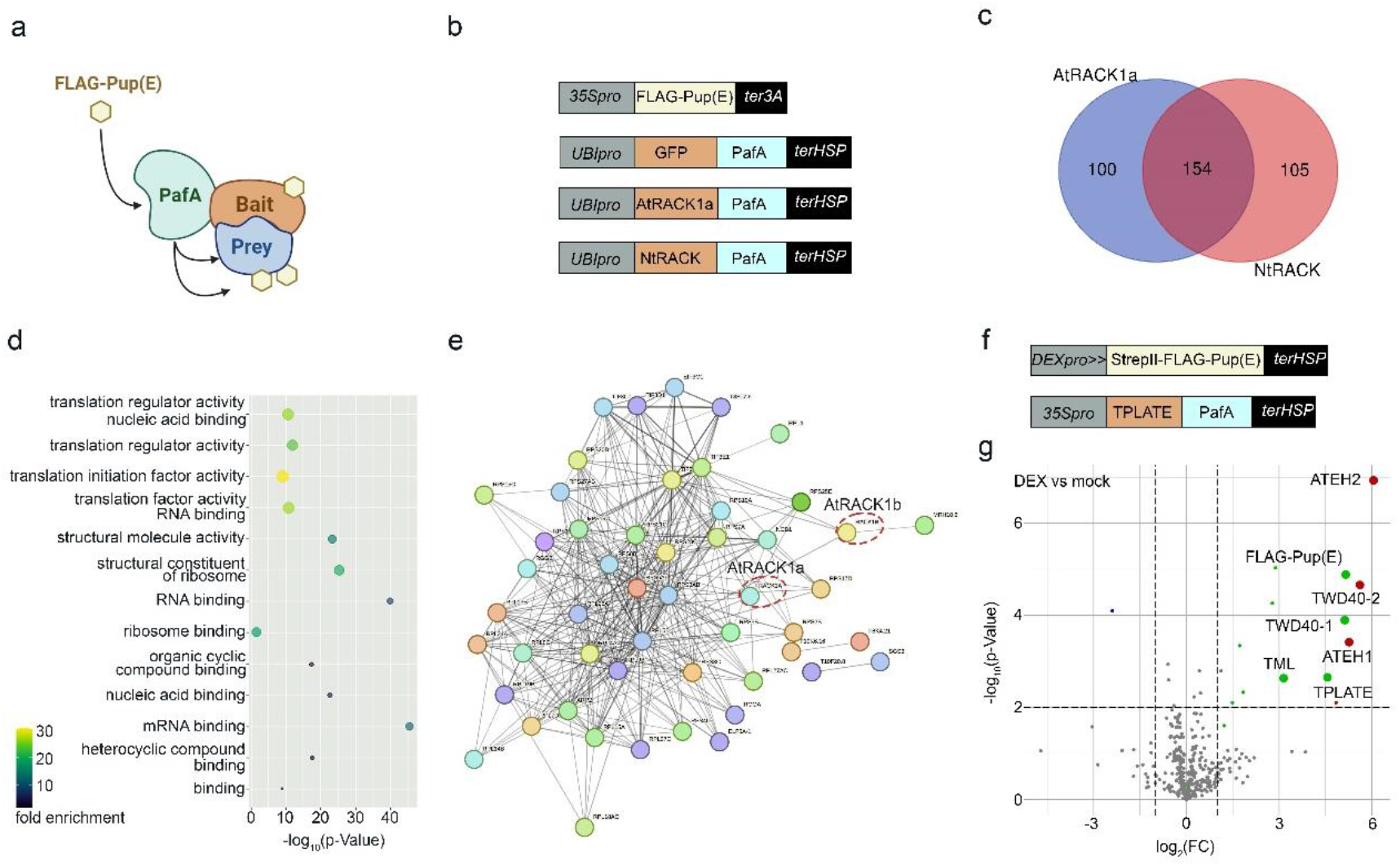
Implementation of PUP-IT in plants. **a**, Schematic image of the PUP-IT proximity labeling system. **b**, Example vectors designed to express the FLAG-Pup(E), and PafA fused to GFP/AtRACK1a/NtRACK. Vector systems contain both the PafA and PUP(E) cassettes on the same backbone. **c**, Overlap of the identified proteins by AtRACK1a and NtRACK as PafA-baits compared to PafA-GFP, respectively, expressed transiently in tobacco leaves. **d**, Enrichment of the molecular function of proteins in the overlapping area of (c). **e**, A PPI network of overlapping proteins between our PafA analyses and proteins indicated as RACK1 interactors in the String database (https://string-db.org/). Nodes indicate proteins and edges indicate interactions, with line thickness indicating the strength of data support in String. **f**, Vectors designed to inducibly express STREPII-FLAG-Pup(E), combined with constitutive expression of PafA fused to TPLATE. **g**, Volcano plots showing protein enrichment in the DEX and mock treatment groups of constructs shown in f expressed in Arabidopsis suspension cells. Through a two-sided t-test (determined by permutation-based FDR calculation, using thresholds FDR = 0.05 and S0 = 1), green spots represent proteins only detected in the DEX group [with relative log2(FC) and -log10(p-Value) compared to the imputation], and red spots represent proteins with high abundance. The Pup(E) and TPLATE complex subunits were highlighted by increased spot size.

The PUP-IT system is based on the enzyme PafA, which catalyses the attachment of a Pup(E) peptide to lysine residues on proteins in close vicinity of the enzyme^4^. The PUP-IT stems from a protein degradation system in prokaryotes that is not present in eukaryotes, and thus Pup(E)-labeling of proteins may be used to enrich and identify them without noticeable background^4^. To implement PUP-IT in plant science, we targeted several proteins and performed experiments in a range of plant systems. We introduced different tags on the Pup(E), e.g. FLAG-Pup(E) and StrepII-FLAG-Pup(E), to readily enrich Pup(E)-Tagged proteins on the same vector backbone as the PafA, which allows implementation of the system via a single transformation event.

We first tested the PUP-IT system using the scaffold protein RACK1 as bait in transient infiltration assays of tobacco leaves. RACK1 has many known interactors^6^, and its activity is mainly associated with protein translation and ribosome function. RACK1 has three paralogs in Arabidopsis, of which RACK1A (AtRACK1a) is the most studied^6^. We therefore selected AtRACK1a and one of the tobacco homologs (Fig. S1a), referred to as NtRACK1, as baits in our PUP-IT approach, with PafA-tagged GFP as control (Fig. 1b). We introduced a flexible linker between the PafA and the RACK1 to minimize interference of the PafA on the native bait function. We based the linker length and content (glycine-serine linkers) on the structure of PafA and RACK1 and Alphafold2 to ensure the flexibility and functionality of the linked proteins (Fig. S1b)^7-10^. We infiltrated the constructs into tobacco leaves, performed subsequent enrichment of FLAG-PUP-enriched proteins and undertook mass-spectrometry (MS) analyses to identify target proteins of the different RACK1s (Fig. S1c-e). We performed binary analyses (screening proteins that are only detected in one group) and significance tests to identify enriched proteins. We found 254 proteins enriched in the AtRACK1a and 259 in the NtRACK1 compared to the GFP control (Fig. 1c) (Dataset S1). Importantly, we found that 154 of the enriched proteins overlapped between the AtRACK1a and NtRACK1 samples (Fig. 1c), corroborating our expectation that the RACK1s in Arabidopsis and tobacco engage with similar proteomes. To assess what processes were enriched among these proteins we next undertook Gene Ontology (GO) analyses and found that terms related to protein translation, RNA binding and ribosome function were among the most enriched terms for the proteins (Fig. 1d). To further substantiate that we captured known interactors of RACK1, we overlaid our PUP-IT identified interactors with what is catalogued in the STRING database (https://string-db.org/). Note that we here mapped tobacco proteins onto the Arabidopsis proteome and thus may miss certain interactors. Figure 1e shows that out of the 154 proteins enriched from our RACK1s experiments, we found that 51 proteins overlapped with a stable RACK1 complex from STRING, and several other interacting proteins formed smaller connected satellite units.

We next tested the PUP-IT approach in another commonly used system for proteomics in plant biology; Arabidopsis PSB-D suspension cells. Here, we chose to investigate the TPLATE complex (TPC), which is a key endocytic protein complex in plant cells containing eight subunits^11^. We, therefore, fused the PafA with TPLATE via a GSL linker and combined it with expression of StrepII-FLAG-Pup(E) under a dexamethasone (DEX)-inducible promoter on the same T-DNA (Fig. 1f). We first optimized the time points and concentrations of DEX treatment by detecting the StrepII-FLAG-Pup(E) labeled TPLATE bait (Fig. S1f-g). Next, we affinity purified the pupylated proteins and probed the beads fraction with an antibody against AtEH1/Pan1, a subunit of the TPC. The co-purification of the AtEH1/Pan1 subunit served as a first proxy for complex incorporation of the tagged bait (Fig. S1h). We also tested whether we could elute the majority of StrepII-FLAG-Pup(E) labeled proteins with 50mM biotin while minimizing Streptactin contamination, which could interfere with mass spectrometry analysis (Fig. S1i-j). Subsequent proteomics analyses showed that PUP-IT has the capacity to specifically isolate the TPLATE complex as various subunits of the TPLATE complex were significantly enriched or even exclusively found in the DEX group (Fig. 1g) (Dataset S2). Furthermore, the absence of the FLAG-PUP(E) peptides in the control group indicates that the DEX-inducible system in PSB-D is not leaky and therefore allowed tight control over the reaction (Fig. 1g).

The above examples demonstrate the utility of PUP-IT as a proximity labelling system in plant biology. We next employed the PUP-IT system to investigate the membrane-based CELLULOSE SYNTHASE (CESA) complex (CSC), which underpins cellulose synthesis in plant cell walls^12^. Here, we chose to fuse the PafA to the N-terminus of COMPANION OF CESA1 (CC1; Fig. S2a), which is one of the central components of the CSC^13^ and transformed the construct either into tobacco leaf cells (transient infiltration) or into *cc1cc2* double mutant Arabidopsis plants (stable transformation). As a control, we used the plasma membrane-localized protein LTI6B (Fig. S2a). We first analyzed the enriched proteins from the tobacco infiltration and found that many of the known CESA complex proteins were enriched in the CC1 samples as compared to the LTI6B (Fig. 2a, Fig. S2b-c) (Dataset S3). For example, beyond CC1, we found tobacco proteins corresponding to the main CESAs that make up the core of the complex (e.g. CESA1), as well as CESA INTERACTING1 (CSI1) that connects the CSC to underlying microtubules^14^. In addition, we found SHOU4L that aids in the trafficking of the CSC, as well as SRF6 from the STRUBBELIG-RECEPTOR family that is involved in response to cellulose deficiency (Fig. 2a) (Dataset S3) ^15,16^.

**Figure 2:**
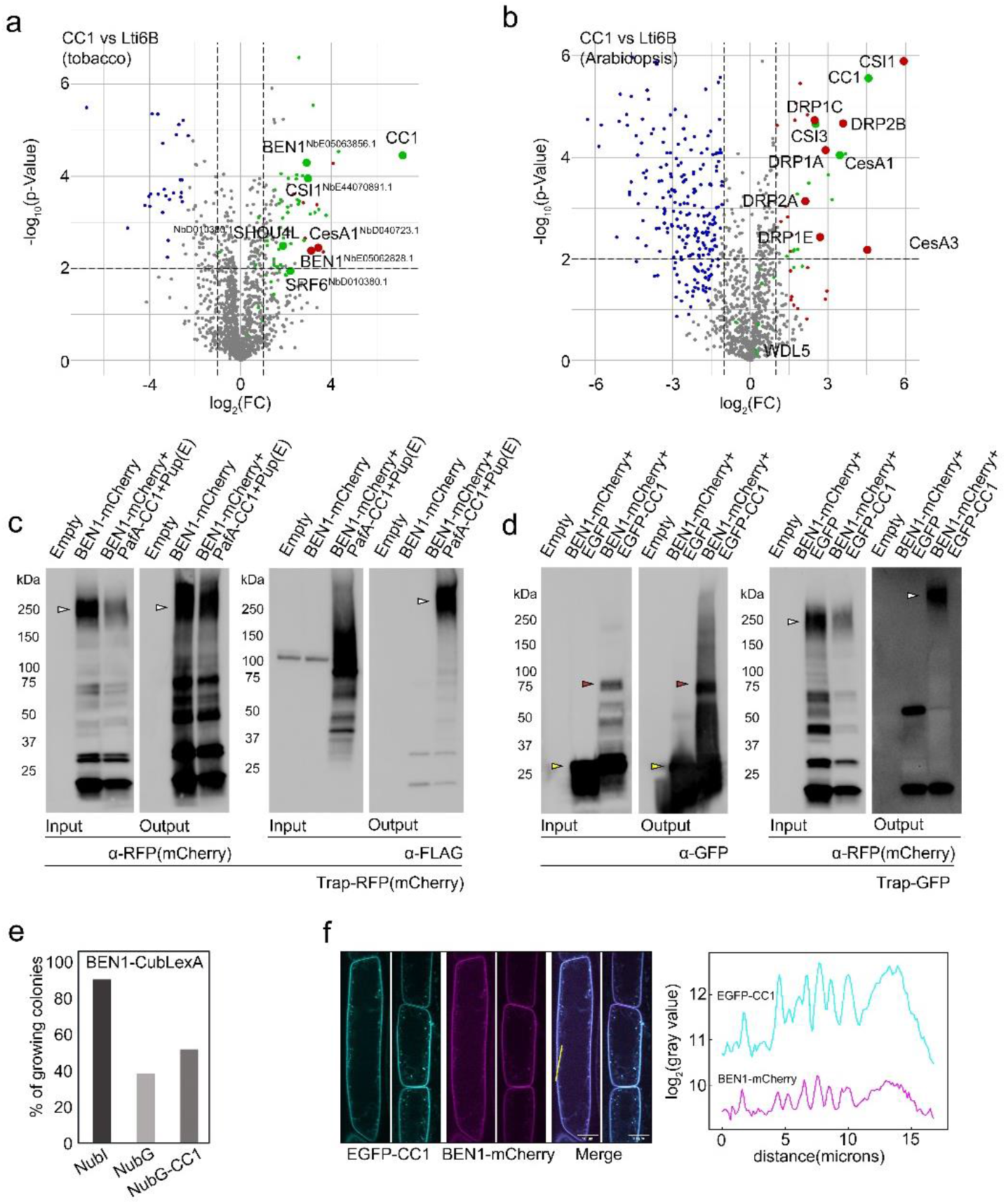
Employment of PUP-IT to identify proteins in cellulose synthesis. **a, b**, Volcano plots showed different enrichment of proteins in the CC1 and Lti6B groups in tobacco leaves (a) and Arabidopsis seedlings (b), respectively, through a two-sided t-test (determined by permutation-based FDR calculation, using thresholds FDR = 0.05 and S0 = 1). Green spots represent proteins only detected in the CC1 group [with relative log2(FC) and - log10(p-Value) compared to the imputation], and red spots represent proteins with high abundance. The proteins related to the CSC and BEN1 were highlighted with increased spot size. **c**, BEN1-mCherry was labelled with FLAG when co-expressed with FLAG-Pup(E) and PafA-CC1 in tobacco leaves. **d**, BEN1-mCherry was co-immunoprecipitated with EGFP-CC1 in tobacco leaves. EGFP was used as control. Arrowheads point to proteins of interest, with the color white corresponding to BEN1-mCherry, yellow corresponding to EGFP and red corresponding to EGFP-CC1. **e**, The membrane split-ubiquitin Y2H was used to detect the interaction between BEN1 and CC1. Approx. 100 colonies were used to quantify interactions and are expressed as ratio of growing vs non-growing colonies. **f**, Co-localization of dual-labelled stable transgenic Arabidopsis lines expressing BEN1-mCherry and EGFP-CC1, Scale bar = 10 μm. The right panel shows fluorescent intensity along transect in the left panel. Seedlings were grown for 5 days, and images were taken in the root elongation zone.

We next investigated the stable CC1 promoter-driven PafA-CC1 Arabidopsis transgenic lines, and Ubiquitin promoter-driven PafA-LTI6B as control. We screened many independent transgenic lines to obtain suitable levels of PafA-LTI6B and PafA-CC1 activity (Fig. S2 d-e). We used six-day-old seedlings as material for the FLAG enrichment, to ensure that we both had sufficient material for enrichment assays and active cellulose synthesis. Here, we again found the components of the core cellulose synthesis machinery, including CESAs (1 & 3) and CSI (1&3)^17^ (Fig. 2b) (Dataset S4). In addition, we found several proteins of the DYNAMIN family that drive CSC internalization^18^. Notably, we also found WDL5, which is involved in microtubule organization during salt stress^19^, similar to that of CC1. We note that despite using different organisms, conditions, transformation approaches and organs, we identified the key core proteins of the CSC using the PafA-CC1 in tobacco and Arabidopsis. However, we also note that the CC1 interactomes vary across the comparison. These data support PUP-IT as an important tool to detect PPIs, and corroborate that the interacting proteome of a given protein may vary depending on development, environment and biological system.

To showcase the effectiveness of PUP-IT in identifying new components of a process, we chose to characterize BEN1/BIG5/AtMIN7’s role in cellulose synthesis (Highly enriched in PaFA-CC1 samples from transient tobacco assays and in both PafA-CC1 and PafA-Lti6B samples from stable Arabidopsis transgenic lines; Fig. 2a, Datasets S3-S4). BEN1 is an ARF-GEF involved in the trafficking of proteins between the TGN and the plasma membrane^20^. We first confirmed that BEN1 could be labelled when co-expressed with the FLAG-Pup(E) and PafA-CC1 in tobacco leaves (Fig. 2c). We then corroborated BEN1 interaction with CC1 via co-immunoprecipitation assays in tobacco leaves (Fig. 2d), and through split-ubiquitin assays in yeast (Fig. 2e). We confirmed that the GFP-CC1 and BEN1-mCherry co-localized at the plasma membrane and TGN in stable Arabidopsis lines (Fig. 2f), corroborating the function of BEN1 at the TGN^20,21^. Given the role of BEN1 in TGN biology, we next assessed how CC1 and BEN1 may connect at this endomembrane by exposing *cc1cc2* mutant seedlings to changes in TGN function using BrefeldinA; a potent ARF-GEF inhibitor that lead to BFA-related endomembrane aggregates in the cytoplasm^22^. Here, the *cc1cc2* mutant seedlings were more sensitive to long-time BFA treatment than wild type, and showed similar phenotype as that of the published T-DNA mutant of BEN1 (Fig. S3a-b). We next generated *ben1cc1cc2* triple mutants to investigate possible genetic interactions of the genes. Notably, *ben1cc1cc2* triple mutant plants displayed reduced rosette leaf expansion, which corresponded to lower levels of cellulose, as compared to the wild-type, *ben1* and *cc1cc2* mutants (Fig. 3a-b; S3c). In addition, the *ben1cc1cc2* triple mutant seedlings displayed increased sensitivity to the cellulose synthesis inhibitor isoxaben (Fig. 3c-d; S3d).

**Figure 3:**
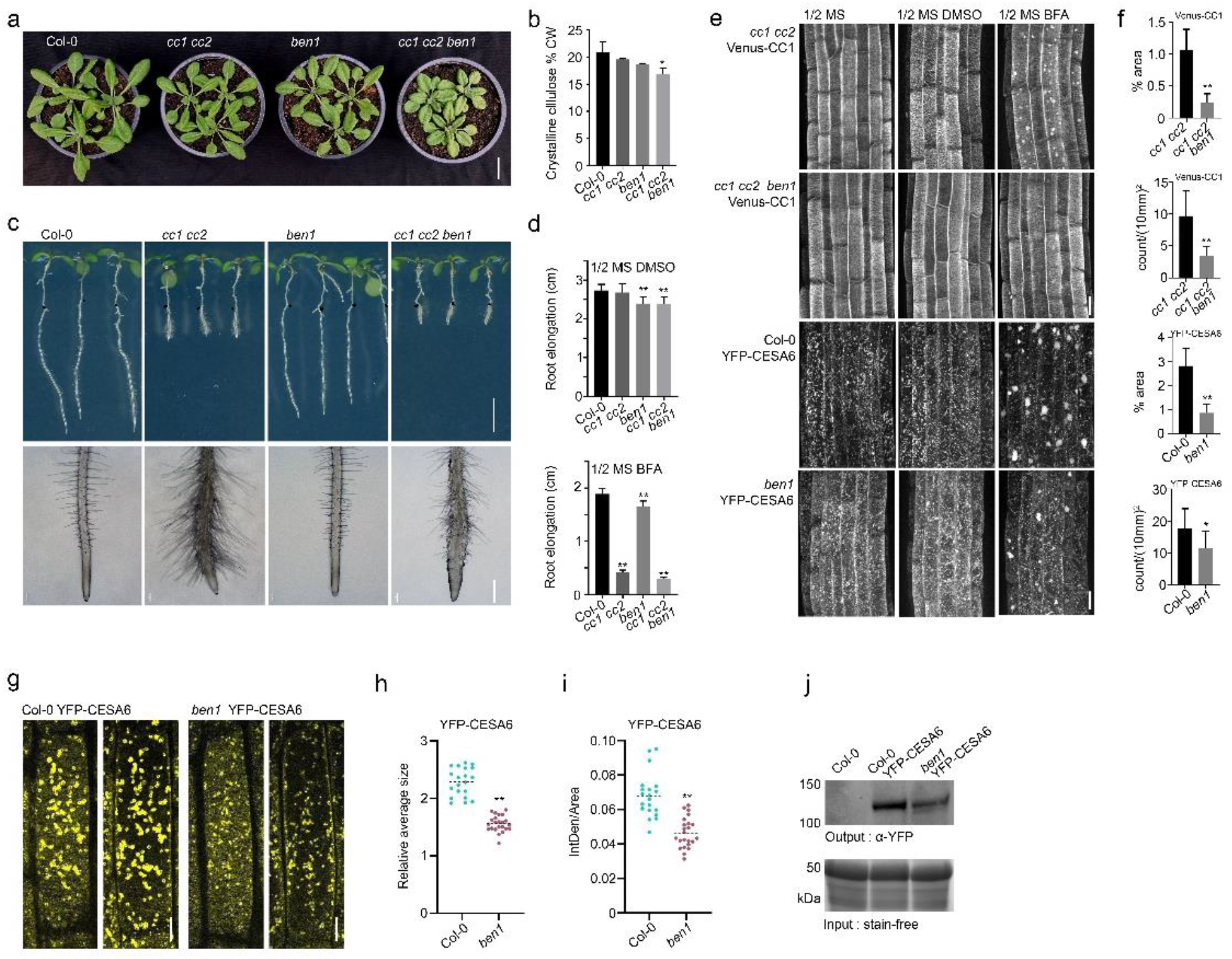
BEN1 impacts trans-Golgi function and cellulose synthesis. **a**, Five-day-old seedlings were transferred to soil and grown for 18 days in the greenhouse, scale bar = 2 cm. **b**, Rosette leaves were harvested from plants growing for 3 weeks in the greenhouse and cellulose was measured according to Methods (Student t-test, n = 3, *P < 0.05). **c, d**, Three-day-old seedlings grown on ½ MS were transferred to ½ MS with 2 nM isoxaben or control for an additional 4 days, scale bar = 0.5 cm (root elongation), scale bar = 0.5 mm (root tip area) (c). Root lengths were measured for seedlings transferred to the different media (Student t-test, n ≥ 10, **P < 0.01) (d). **e, f**, Venus-CC1 and YFP-CESA6 in *ben1* or control backgrounds were imaged following BFA treatment (50 μM for 30 min), scale bar = 10 μm (e), and BFA body distribution either across the root area using percentage area (% area; upper panel) or count per mm^2^ (lower panel) were used for statistical analyses (Student t-test, n ≥ 6, **P < 0.01 & *P < 0.05). (f). **g-i**, YFP-CESA6 distribution in Col-0 and *ben1* background, max intensity projection from two Z stacks with three steps of 0.3 μm (left panels Z stack starting at the plasma membrane and right panels starting 0.9 μm below the plasma membrane per genotype), scale bar = 10 μm (g), quantification of YFP-CESA6 fluorescence from Golgi bodies with relative size (h) and IntDen (the product of Area and Mean Gray Value)/Area of fluorescence (i), (Student t-test, n ≥ 20, **P < 0.01). **j**, Western blot to detect YFP-CESA6 in Col-0 and *ben1* background using α-YFP(Venus) (Merck, MABE1906). YFP-CESA6 was immunoprecipitated by GFP-Trap Agarose (Chromotek, gta) from 0.5g seedlings grown for 2 weeks on ½ MS plates.

To directly visualize the behaviour of CC1 and the CSC in the *ben1* mutant, we generated Venus-CC1*ben1* (Fig. S3e-f) and YFP-CESA6*ben1* plants. Previous studies showed that mutations in BEN1 affect trafficking of, for example, PIN-FORMED (PIN) and BRASSINOSTEROID INSENSITIVE1 (BRI1), which led to reduced accumulation of the proteins in BFA bodies^20,23^. Consistent with this, we observed reduced accumulation of both Venus-CC1 and YFP-CESA6 in BFA bodies in the *ben1* mutant compared to wild type (Fig. 3e-f). Interestingly, we found that the distribution of YFP-CESA6 fluorescence in the Golgi differed between wild-type and *ben1* cells. Here, the YFP-CESA6 fluorescence was dimmer in the *ben1* mutant and the fluorescence in the Golgi was less intense with narrower Golgi diameter compared to that of wild type (Fig. 3g-i). In addition, the YFP-CESA6 protein levels were reduced in *ben1* as compared to wild type (Fig. 3j). These data indicate that BEN1/BIG5 regulates the trafficking of CC1 and CESAs between the TGN and plasma membrane and that defects in the BEN1/BIG5 lead to reduced cellulose synthesis possibly through a feedback-related decrease in CSC levels.

Protein interactions is an essential tool in biology to understand how proteins work in context to each other and to infer protein function. The implementation of new tools to study PPIs is therefore of substantial interest across all aspects of cell biology. We provide a new tool box for PPI inferences in plant biology and showcase how this tool may be used to identify new components of an essential process in plants. We envision that PUP-IT may become a suitable approach to complement other types of PPI inferences, such as affinity-based purification and biotin-based proximity labelling.

## Methods

### Plant materials and growth conditions

T-DNA insertional lines for *ben1* (SALK_013761C) were obtained from NASC (http://Arabidopsis.info/; ref); the double mutant *cc1 cc2* (SAIL_838_F07, Gabi_654A12) and marker lines Venus-CC1 (*cc1 cc2*), YFP-CesA6 (Col-0)^13^ were from Persson’s lab. The double *cc1 cc2* was crossed to *ben1* and the triple mutant *cc1 cc2 ben1* was identified by *cc1, cc2*, and *ben1* genotyping primers from the F2 generation. Marker line YFP-CESA6 (Col-0) was crossed to *ben1*, and *the* YFP-CESA6 (*ben1*) was identified by *ben1* genotyping primers and imaging of the YFP-CESA6. Marker line Venus-CC1 (*cc1 cc2*) was crossed to the triple mutant *cc1 cc2 ben1*, and the Venus-CC1 (*cc1 cc2 ben1*) was identified by *ben1* genotyping primers and imaging of the Venus-CC1. The newly generated marker lines were examined in the F3 generation to confirm no segregations of the fluorescent markers. Primers used to screen for homozygous insertion lines are present in Table S1.

Arabidopsis seedlings were grown in a growth chamber under long-day conditions (16 h light / 21 °C and 8 h dark / 19 °C) on solid or liquid half mass spectrometry (MS) media with 1% sucrose. After that, seedlings were transferred to soil pots in a greenhouse under set conditions (16 h light / 21 °C and 8 h dark / 19 °C).

### Plasmid construction

Primers used for constructing all the constructs in this manuscript are present in Table S1. Backbones and fragments were assembled with NEBuilder® HiFi DNA assembly mix (New England Biolabs) / In-fusion Master mix (Takara) or ligated using T4 ligase (Thermo Fisher Scientific/New England Biolabs) in corresponding clones.

To create the vectors that express the FLAG-Pup(E) and PafA fused proteins. The fragments of FLAG-Pup(E) + ter3A were first assembled into the AscI site of pMDC32^24^ to produce the destination vector pMDC32_35S::FLAG-Pup(E)*::ter3A::attR1-attR2::terNOS*. The UBI10 promoter and PafA were assembled into a pENTR vector. Then CDS of LTI6B and the pENTR backbone with *UBI10pro::*PafA were amplified and assembled. After that, an LR reaction was performed to integrate the linearized pENTR *UBI10pro::*PafA-LTI6B into the destination vector to produce the pMDC32_35S::FLAG-Pup(E)*::ter3A_+_UBIpro::*PafA-LTI6B*::terNOS*.

pENTR/TOPO modified with multiple cloning sites was used as the backbone for the GFP/AtRACK1a/NtRACK1 vectors. pENTR/TOPO *CC1pro::*mNeonGreen-CC1*::terHSP* was used as the backbone for the CC1 vectors. Then, the HpaI or ApaLI (New England Biolabs) linearized pENTR of attL1-*UBIpro::*GFP/AtRACK1a/NtRACK1-PafA::*terHSP-*attL2 and attL1-*UBIpro/CC1pro::*PafA-CC1::*terHSP-*attL2 were integrated into *pMDC32*_35S::FLAG-Pup(E)*::ter3A::attR1-attR2::terNOS* respectively through LR reactions.

To create the pGGK-35S::GRLhG4>>A-G, vector pGG-A-35S-B, pGGD-linker-E, pGG-E-35St-F, pGG-F-AarI_linker-G^25^ and the pSW610 (pGG-B-GR-LhG4-D^26^ ; addgene #115992) were assembled into a pGGK A-G vector via a Golden Gate reaction as previously described^25,27^. After AarI (Thermo Scientific™) digestion, an A-ccdB/CmR-G PCR fragment was inserted into the resulting vector via ligation using T4 DNA ligase to generate pGGK-35S::GRLhG4>>A-ccdB-G. pGGK-35S::GRLhG4>>p6xOP::StrepII-FLAG-PUP*::tHSP18*.*2*_+_35S::TPLATE-GSL-PafA*::tHSP18*.2 was created with two rounds of iterative Golden Gate assembly as previously described^27^. In the first Golden Gate reaction pGGK-35S::GRLhG4>>A-ccdB-G was combined with pSW180a - pOp6 (pGG-A-p6xOP-B)^26^, pGGB-StrepII-C, pGGC-FLAG-PUP-D, pGG-D-Decoy_v2-E (D017), pGG-E-tHSP18.2M-F^28^ and pGG-F-A-AarI-SacB-AarI-G-G^25^. After AarI digestion, the resulting vector was combined with pGGA-35S-C^29^, pGGC-TPLATE-D, pGGD-GSL-PafA-E, pGG-E-tHSP18.2M-F and pGG-F-linkerII-G^25^ in a second Golden Gate reaction.

### PUP-IT proteins expression in tobacco, Arabidopsis thaliana PSB-D cell and stable transgenics

The Agrobacterium gv3101 transformed with pMDC32_35S::FLAG-Pup(E)*::ter3A* _+_*UBIpro::*GFP/AtRACK1A/NtRACK1-PafA*::terHSP* and _+_*UBIpro::* PafA-Lti6B*::terNOS*/CC1*::terHSP* respectively were cultured and infiltered into tobacco (*Nicotiana benthamiana*) leaves along with strain P19^30^ (OD600 = 0.5 in infiltration buffer) as previously described^31^. The tissues were collected 2 days after plants grown in the greenhouse.

The pGGK-35S::GRLhG4>>p6xOP::StrepII-FLAG-PUP*::tHSP18*.*2*_+_35S::TPLATE-GSL-PafA*::tHSP18*.*2* construct was transformed into dark grown PSB-D suspension culture cells as described before^32^. The cultures were harvested 3 days after subculturing. StrepII-FLAG-PUP(E) expression was induced with 1, 10, 50, 100 or 200 μM dexamethasone as indicated (stock solution: 50mMol Dexamethasone in DMSO) for 1 or 24 hours as indicated. DMSO was used as a mock treatment.

The Agrobacterium gv3101 transformed with pMDC32_35S::FLAG-Pup(E)*::ter3A* _+_*UBIpro::*PafA-Lti6B*::terNOS*/ *CC1pro::*PafA-CC1*::terHSP* respectively were used to produce Arabidopsis transgenics through flower dipping^33^. After hygromycin selection, positive transgenics were identified by western blot with FLAG antibody. Seedlings were grown in liquid half MS for five days in culture bottle to generate tissues for the followed IP-MS.

### SDS-PAGE and Western blot

Samples from Tobacco leaves and Arabidopsis transgenics in 1× Laemmli loading buffer (BioRad) with reducing agent (100 mM DTT) were boiled for 10 min at 95 °C. Protein were loaded on a 12% or 4-15% SDS–PAGE TGX gel (BioRad) and subsequently blotted on a polyvinylidenedifluoride membrane (BioRad). Membranes were blocked in 5% skimmed milk overnight at 4°C on a shaking device. Next, the blots were incubated with primary antibodies α-FLAG (Merck, F3165, 1:2000), α-GFP (Chromotek, 3H9, 1:2000), α-YFP (Venus) (Merck, MABE1906, 1:1000) and secondary antibodies α-mouse-HRP (Agilent, P0260, 1:5000), α-rat-HRP (GE HealthCare, NA935V, 1:5000) in 5% skimmed milk for 1h at room temperature on a shaking device. For RFP (mCherry) detection, membranes were blocked for 2 h at room temperature, incubated with primary antibody α-RFP (mCherry) (Chromotek, 6G6, 1:2000) overnight at 4°C, and secondary antibody for 1 h at room temperature on a shaking device.

Samples from PSB-D suspension culture cells in 1× Laemmli loading buffer (BioRad) with reducing agent (1× NuPage [Invitrogen] or 100 mM DTT) were boiled for 10 min at 95 °C. Protein were loaded on a 4–20% SDS–PAGE TGX gel (BioRad) and subsequently blotted on a polyvinylidenedifluoride membrane (BioRad). Membranes were blocked in 5% skimmed milk overnight at 4°C on a shaking device. Next, the blots were incubated with primary antibodies α-FLAG (Merck, F1804, 1:1000], α-TPLATE^34^(1:1000), α-AtEH1/Pan1^35^(1:2000) and secondary antibodies α-mouse-HRP (Cytiva, NA931, 1:10,000), α-rabbit-HRP (Cytiva,NA934, 1:10,000) in 3% skimmed milk for 1h at room temperature on a shaking device.

### Protein extraction and Pull-Down

Tissues from tobacco leaves (2-3 g) were sampled and ground in liquid nitrogen to a fine powder. Total proteins were extracted with extraction buffer (1:3 [W/V]-material: buffer), [20 mM Tris-HCl, pH 7.4, 150 mM NaCl, 5 mM MgCl2, 1 mM EDTA, 1% Triton X-100, and 0.1% protease inhibitor cocktail (Roche)] for 30 minutes on ice, and centrifuged at 4°C at 12,000 rpm for 30 min. The supernatant was incubated with equilibrated anti-Flag antibody coupled magnetic beads (Thermo Scientific™, A36797) for 2 hours at 4°C. Beads were washed three times with extraction buffer [20 mM Tris-HCl, pH 7.4, 150 mM NaCl, 5 mM MgCl2, 1 mM EDTA, 1% Triton X-100 and 0.1% protease inhibitor cocktail (Roche)], then washed three times with wash buffer [20 mM Tris-HCl, pH 7.4, 150 mM NaCl, 5 mM MgCl2, 1 mM EDTA and 0.1% protease inhibitor cocktail (Roche)] and three times with 1 X PBS. After that the beads were stored at −80°C until LC-MS/MS analysis.

A similar process was performed on tissues from Arabidopsis seedlings (1-2 g). The extraction buffer was replaced by [50 mM HEPES, pH 7.5, 150 mM NaCl, 10 mM NaF, 5 mM EDTA, 1% Triton X-100, 1% PVP and 0.1% protease inhibitor cocktail (Roche)], and the wash buffer was replaced by [50 mM HEPES, pH 7.5, 150 mM NaCl, 10 mM NaF, 5 mM EDTA, 1% PVP and 0.1% protease inhibitor cocktail (Roche)].

Total protein extracts from liquid N2 ground PSB-D cells (3 g/replicate) were generated by adding extraction buffer (2:3 - buffer:material), (25mM TRIS-HCl pH7.6, 15mM MgCl2, 150mM NaCl, 15mM pNO2phenylPO4, 60mM B-glycerophosphate, 0.1mM Na3VO4, 1 mM NaF, 1mM PMSF, 1um E64, 0.5mM EDTA, 0.50% NP40, 5%ethyleneglycol, 0.1% protease inhibitor cocktail EDTA-free (Roche)]) and incubating for 1 hour at 4°C. Subsequently, the lysate was cleared by 2 centrifugation steps at 20000g at 4°C. Then the lysate was incubated with equilibrated Strep-Tactin®XT 4Flow® high-capacity resin (100ul beads/ replicate) for 2 hours at 4°C. Next, the beads were washed 3 times (10 column volumes total). Bound proteins were eluted using 2x50ul 50mM biotin in 50mM NH4CO3. Subsequently, the supernatant was cleared by 1minute centrifugation at 20000g. Protein samples eluted with biotin were digested in-solution with 1 ug Trypsin/LysC (Promega) overnight at 37°C, followed with an additional 0.5ug Trypsin/LysC for 2 hours. Finally, digested proteins were cleaned up with C-18 Omix tips (Agilent). Digests containing the cleaved peptides were dried in a SpeedVac and stored at −20°C until LC-MS/MS analysis.

### Mass spectrometry and data analysis

Washed beads were incubated for 30 min with elution buffer 1 (2 M Urea, 50 mM Tris-HCl pH 7.5, 2 mM DTT, 20 μg/ml trypsin) followed by a second elution for 5 min with elution buffer 2 (2 M Urea, 50 mM Tris-HCl pH 7.5, 10 mM Chloroacetamide). Both eluates were combined and further incubated at room temperature overnight. Tryptic peptide mixtures were acidified to 1% TFA and loaded on Evotips (Evosep). Peptides were separated on 15 cm, 150 μM ID columns packed with C18 beads (1.9 μm) (Pepsep) on an Evosep ONE HPLC applying the ‘30 samples per day’ method, and injected via a CaptiveSpray source and ten μm emitter into a timsTOF pro mass spectrometer (Bruker) ran in PASEF mode^36^.

Peptides were re-dissolved in 20 μl loading solvent A (0.1% trifluoroacetic acid in water/acetonitrile (ACN) (98:2, v/v)) of which 2 μl was injected for LC-MS/MS analysis on an Ultimate 3000 RSLCnano system in-line connected to a Q Exactive HF Biopharma mass spectrometer (Thermo). Trapping was performed at 20 μl/min for 2 min in loading solvent A on a 5 mm trapping column (Thermo scientific, 300 μm internal diameter (I.D.), 5 μm beads). 250 mm Aurora Ultimate, 1.7μm C18, 75 μm inner diameter (Ionopticks) kept at a constant temperature of 45°C. Peptides were eluted by a non-linear gradient starting at 1% MS solvent B reaching 26% MS solvent B (0.1% FA in acetonitrile) in 30 min, 44% MS solvent B in 38 min followed by a 5-minute wash at 56% MS solvent B starting at 40 minutes and re-equilibration with MS solvent A (0.1% FA in water) at a flow rate of 300 nl/min.

The mass spectrometer was operated in data-dependent mode and raw files were processed using the MaxQuant software (version 2.0.3.0)^37^. Peak lists were searched against the proteome of released *Nicotiana benthamiana*^38^ or the proteome of Arabidopsis (Uniprot: UP000006548_3072 and UP000006548_3702_additional) combined with 262 common contaminants by the integrated Andromeda search engine. Additionally, the FLAG-Pup(E) sequence and other commonly used tag sequences were added. Mass tolerance on precursor ions was set to 4.5 ppm and on fragment ions to 20 ppm. Match between runs and MS1-based Label Free Quantification (LFQ) were on. The false discovery rate was 1 % for both peptides (minimum length of 7 amino acids) and proteins.

The protein groups result file was uploaded in Perseus (2.0.10.0)^37^. Reverse hits, contaminants and only identified by site identifications were removed. LFQ intensity values were log2 transformed. For each comparison between experiment and control group, identifications were filtered for 50% values in total mimicking the assumption that specific proteins would be only detected in one group. A binary analyse was first performed to screen proteins as unique that were detected in all replicates within an experiment group but no detection in the corresponded control. For relative comparation, missing values were imputated with values around the detection limit, randomly drawn from a normal distribution with a width equal to 0.3 and a downshift equal to 1.8. Then a two-sided t-test was performed with the filter of log2(FC) > 1, -log10(p-value) > 2 and corrected false positive by permutation-based FDR calculation, using thresholds FDR = 0.05 and S0 = 1. The significant candidate lists were generated through collection from proteins unique and relatively high abundance through t-test, then plotted in volcano summary by R.

Proteomics data are available from PRIDE: PXD048346

### Plant phenotyping

Seedlings of wild type (Col-0) and corresponding mutants were grown on half MS plates for 3 days, then transferred to the plates with 5 μM BFA for 7days or 2nM Isoxaben for 4 days (DMSO as control). The elongated root length was measured by FIJI-ImageJ and root tip were captured by (KEYENCE, VHX-7000).

7 days seedlings were transferred into soil for 20 days in the greenhouse and pictured to record the growth of rosette leaves. 3 samples were harvested for each group, and each sample contained leaves from two pots. The crystalline cellulose quantification was conducted as described^39,40^.

### Immunoprecipitation and split yeast two hybrid assay

The coding sequences of *CC1* and *BEN1* (without stop codon) were amplified by PCR and subcloned into the entry vectors PUCL3L2 (Plasmid #114019, addgene) and PUCL1L4 (Plasmid #114018, addgene) respectively. Then BEN1, CC1 and BEN1 were introduced into the pFRETgc-2in1-NC (Plasmid #105121, addgene) vector by LR reaction to generate the pFRETgc 35S::EGFP + 35S::BEN1-mCherry and pFRETgc 35S:: EGFP-CC1 + 35S::BEN1-mCherry. pFRETgc 35S::EGFP + 35S::BEN1-mCherry were co-expressed with pMDC32_35S::FLAG-Pup(E) _+_UBIpro::::PafA-CC1 or not in tobacco leaves, then proteins were immunoprecipitated by RFP-Trap Agarose (Chromotek, rta). pFRETgc 35S::EGFP + 35S::BEN1-mCherry and pFRETgc 35S:: EGFP-CC1 + 35S::BEN1-mCherry were expressed in tobacco leaves respectively, then proteins were immunoprecipitated by GFP-Trap Agarose (Chromotek, gta).

The split yeast two hybrid assay was performed by employing the approach as developed^41^. Briefly, the CDS of CC1 and BEN1(without stop codon) were subcloned in the pENTR/DTOPO then introduced into the destination vectors pNX32-DEST (Plasmid #105083, addgene) and pMetYC-DEST (Plasmid ##105081, addgene) through LR reaction. The pNX32:CC1 and pMetYC:BEN1 were co-transformed into the yeast strain NMY51. The transformed yeast strains were plated in SD/-Leu/-Trp or SD/-Leu/-Trp/-His/-Ade/+0.3mM Met medium and placed in a 30°C constant temperature incubator for 3-5 days for selections, 96 colonies from each combination were randomly picked for the survival rate calculation.

### Drug treatments

Seedlings were grown on half MS plates vertically for 4 days, then submerged in 3 ml of solution containing the chemical agent(s) in 6-well cell culture plates and incubated in the growth chamber for various time points before imaging. Brefeldin A was dissolved in dimethyl sulfoxide (DMSO). Working solutions of chemicals were freshly diluted in liquid half MS media from stocks immediately before use.

### Fluorescence imaging and analysis

The pictures were taken by Confocal Microscopes 3i CSU-W SoRa Spinning Disk and processed by Fiji for the followed analysis. The vector pFRETgc 35S:: EGFP-CC1 + 35S::BEN1-mCherry was transformed into the Col-0 and selected by BASTA for positive transgenics. Images were taken to collect signals of EGFP (C1: 488/563, 30%, 100 ms) and mCherry (C2: 563/488, 30%, 200 ms), with drawing line defined region of interest to produce intensity plot of co-efficient subcellular localization. For BFA treatment, Venus-CC1 and YFP-CESA6 were excited by laser 515 nm (30%, 200 ms), and observed under Len 60 X with z-step of 0.3 μm. For the YFP-CESA6 distribution in Golgi, images were taken under Len 100 X with settings above. Z-projects with maximum intensity from 63 steps were produced to view the BFA body distributions. Z-projects with maximum intensity from 3 steps were produced to view the YFP-CESA6. Particle Analysis in Fiji were employed to calculate the signals reflecting BFA body and Golgi.

## Acknowledgements

We acknowledge Prof Jürgen Kleine-Vehn for codon optimised templates of PafA and Pup(E). We would like to thank Liu Wang for sharing unpublished marker lines Venus-CC1 (*cc1 cc2*). We thank Michael Kraus, Moritz Nowack, and Norbert Bollier for sharing unpublished golden gate building blocks. We thank Michaël Vandorpe for assisting with the golden gate cloning expression clones. We thank Geert De Jaeger and Jelle Van Leene for constructive discussions and the Interactomics Facility of the Center for Plant Systems Biology (VIB) for assisting with PSB-D cell culture transformation. We thank the Proteomics Research Infrastructure (PRI) at the University of Copenhagen (UCPH) supported by NNF19SA0059305 and the VIB proteomics core for the mass spectrometry data collection. We thank the Center for Advanced Bioimaging (CAB) at the UCPH for the imaging support. A.D.M. is supported by a PhD grant (1124621N) from the Research Foundation– Flanders (FWO). G.A.K. is funded by a DECRA Fellowship from the Australian Research Council (DE210101200). S.P. was funded by a Villum, two Novo Nordisk, and Danish National Research Foundation grants (25915, 19OC0056076, 20OC0060564, DNRF155, respectively).

## Supplementary information

**Figure S1: RACK1 and TPLATE pupylation**.

**a**, A phylogenetic tree comparing the RACK proteins from tobacco and Arabidopsis. **b**, Glycine-Serine Linkers (GGGGSGGG) was introduced between the PafA and AtRACK1a with structure predicted by Alphafold2 (https://alphafold.ebi.ac.uk/). **c**, Western blot to detect the FLAG-Pup(E) labelled proteins from RACK1 transient expression in tobacco leaves. **d, e**, Pulldown of FLAG-Pup(E) labelled proteins. Coomassie blue stain indicated the total proteins in lanes (c) and the same amount of sample was loaded to detect the FLAG-Pup(E) labelled proteins with FLAG antibody (d). **f**, Inducible expression of StrepII-FLAG-Pup(E) after one hour and after 24 hours of 50μM dexamethasone (DEX) treatment in PSB-D cells constitutively expressing TPLATE-GSL-PafA, visualized using WB detection of Pup(E) via anti-FLAG and TPLATE via anti-TPLATE. The open arrowhead points to the endogenous TPLATE, while the closed arrowhead indicates the tagged form with PafA. Note that the band of the tagged form becomes blurry after 24h of induction, suggestive of post-translational modification with Pup(E). **g**, expression analysis of StrepII-FLAG-Pup(E) in the presence of TPLATE-GSL-PafA following a 24-hour treatment with different concentrations of dexamethasone (DEX). The 10μM concentration yielded the strongest pupylation level and was retained for further experiments. The open and closed arrowheads indicate the endogenous TPLATE as well as the tagged form. **h**, Pulldown experiment of StrepII-tagged proteins in PSB-D cells expressing TPLATE-GSL-PafA following a mock or 50μM dexamethasone treatment to induce expression of StrepII-FLAG-Pup(E). Next to the anti-FLAG detection, we also detected AtEH1/Pan1 as a proxy for complex incorporation of the PafA-tagged bait. **i j**, Pulldown experiment of StrepII-tagged proteins in PSB-D cells expressing TPLATE-GSL-PafA following a mock or a 10μM dexamethasone treatment (DEX) to induce expression of StrepII-FLAG-Pup(E), StrepII-tagged proteins (asterisk) were eluted from the beads using 50mM biotin. Strep-TactinXT® (black arrowhead) remains in the bead fraction and does not visibly contaminate the eluate.

**Figure S2: CC1 enrichment via pupylation**.

**a**, Vector design to express the FLAG-Pup(E), and PafA fused LTI6B/CC1.**b**, Western blot to detect the FLAG-Pup(E) labelled proteins from transient expression in tobacco leaves. Western blot was probed with anti-FLAG antibody. **c**, Pulldown of FLAG-Pup(E) labelled proteins from tobacco leaves and detection of the proteins using FLAG antibody. **d e**, Western blot to screen transgenic Arabidopsis that expressed the PUP-IT system with LTI6B (d) and CC1 (e). **f**, Pulldown of FLAG-Pup(E) labelled proteins in Arabidopsis, and subsequent detection of pupylated proteins using a FLAG antibody.

**Figure S3: BEN1 affects trans-Golgi function and cellulose synthesis**.

**a**, Col-0 and CC1, BEN1 transgenic lines grown on media with DMSO (control) or BFA for seven days. Scale bar = 1 cm (a). **b**, Root length measurements from seedlings as those in a (Student t-test, n ≥ 10, **P < 0.01). **c**, Genotyping to identify the *cc1 cc2 ben1* triple mutant, scale bar = 2 cm. **d, e**, Col-0 and CC1, BEN1 transgenic lines grown on DMSO (control) or isoxaben supplemented media. Scale bar = 0.5 cm. **f**, Root length measurements from seedlings as those in e (Student t-test, n ≥ 10, **P < 0.01).

**Table S1. Primers used in this study**.

**Dataset S1. Proteomics data for AtRACK1a and NtRACK1**.

**Dataset S2. Proteomics data for TPLATE**.

**Dataset S3. Proteomics data for CC1 in tobacco**.

**Dataset S4. Proteomics data for CC1 in Arabidopsis**.

## References

1 Xing, S., Wallmeroth, N., Berendzen, K. W. & Grefen, C. Techniques for the Analysis of Protein-Protein Interactions in Vivo. Plant Physiol 171, 727–758, doi:10.1104/pp.16.00470 (2016).

2 Bosch, J. A., Chen, C. L. & Perrimon, N. Proximity-dependent labeling methods for proteomic profiling in living cells: An update. Wiley Interdiscip Rev Dev Biol 10, e392, doi:10.1002/wdev.392 (2021).

3 Yang, X. et al. Proximity labeling: an emerging tool for probing in planta molecular interactions. Plant Commun 2, 100137, doi:10.1016/j.xplc.2020.100137 (2021).

4 Liu, Q. et al. A proximity-tagging system to identify membrane protein-protein interactions. Nat Methods 15, 715–722, doi:10.1038/s41592-018-0100-5 (2018).

5 Ye, R. Q. et al. Glucose-driven TOR-FIE-PRC2 signalling controls plant development. Nature 609, 986–+, doi:10.1038/s41586-022-05171-5 (2022).

6 Islas-Flores, T., Rahman, A., Ullah, H. & Villanueva, M. A. The Receptor for Activated C Kinase in Plant Signaling: Tale of a Promiscuous Little Molecule. Front Plant Sci 6, 1090, doi:10.3389/fpls.2015.01090 (2015).

7 van Rosmalen, M., Krom, M. & Merkx, M. Tuning the Flexibility of Glycine-Serine Linkers To Allow Rational Design of Multidomain Proteins. Biochemistry 56, 6565–6574, doi:10.1021/acs.biochem.7b00902 (2017).

8 Jumper, J. et al. Highly accurate protein structure prediction with AlphaFold. Nature 596, 583–+, doi:10.1038/s41586-021-03819-2 (2021).

9 Ozcelik, D. et al. Structures of Pup ligase PafA and depupylase Dop from the prokaryotic ubiquitin-like modification pathway. Nat Commun 3, 1014, doi:10.1038/ncomms2009 (2012).

10 Ullah, H. et al. Structure of a signal transduction regulator, RACK1, from. Protein Sci 17, 1771–1780, doi:10.1110/ps.035121.108 (2008).

11 Gadeyne, A. et al. The TPLATE Adaptor Complex Drives Clathrin-Mediated Endocytosis in Plants. Cell 156, 691–704, doi:10.1016/j.cell.2014.01.039 (2014).

12 Pedersen, G. B., Blaschek, L., Frandsen, K. E. H., Noack, L. C. & Persson, S. Cellulose synthesis in land plants. Mol Plant 16, 1228, doi:10.1016/j.molp.2023.05.008 (2023).

13 Endler, A. et al. A Mechanism for Sustained Cellulose Synthesis during Salt Stress. Cell 162, 1353–1364, doi:10.1016/j.cell.2015.08.028 (2015).

14 Liu, Z. Y. et al. Cellulose-Microtubule Uncoupling Proteins Prevent Lateral Displacement of Microtubules during Cellulose Synthesis in. Dev Cell 38, 305–315, doi:10.1016/j.devcel.2016.06.032 (2016).

15 Polko, J. K. et al. SHOU4 Proteins Regulate Trafficking of Cellulose Synthase Complexes to the Plasma Membrane. Curr Biol 28, 3174–3182 e3176, doi:10.1016/j.cub.2018.07.076 (2018).

16 Chaudhary, A. et al. The Arabidopsis receptor kinase STRUBBELIG regulates the response to cellulose deficiency. Plos Genet 16, doi:ARTN e1008433 10.1371/journal.pgen.1008433 (2020).

17 Lei, L., Li, S. D., Du, J., Bashline, L. & Gu, Y. CELLULOSE SYNTHASE INTERACTIVE3 Regulates Cellulose Biosynthesis in Both a Microtubule-Dependent and Microtubule-Independent Manner in. Plant Cell 25, 4912–4923, doi:10.1105/tpc.113.116715 (2013).

18 Sanchez-Rodriguez, C. et al. The Cellulose Synthases Are Cargo of the TPLATE Adaptor Complex. Mol Plant 11, 346–349, doi:10.1016/j.molp.2017.11.012 (2018).

19 Dou, L., He, K., Higaki, T., Wang, X. & Mao, T. Ethylene Signaling Modulates Cortical Microtubule Reassembly in Response to Salt Stress. Plant Physiol 176, 2071–2081, doi:10.1104/pp.17.01124 (2018).

20 Tanaka, H., Kitakura, S., De Rycke, R., De Groodt, R. & Friml, J. Fluorescence imaging-based screen identifies ARF GEF component of early endosomal trafficking. Curr Biol 19, 391–397, doi:10.1016/j.cub.2009.01.057 (2009).

21 Nomura, K. et al. Effector-triggered immunity blocks pathogen degradation of an immunity-associated vesicle traffic regulator in. P Natl Acad Sci USA 108, 10774–10779, doi:10.1073/pnas.1103338108 (2011).

22 Chardin, P. & McCormick, F. Brefeldin A: The advantage of being uncompetitive. Cell 97, 153–155, doi:Doi 10.1016/S0092-8674(00)80724-2 (1999).

23 Xue, S. et al. Involvement of BIG5 and BIG3 in BRI1 Trafficking Reveals Diverse Functions of BIG-subfamily ARF-GEFs in Plant Growth and Gravitropism. Int J Mol Sci 20, doi:ARTN 233910.3390/ijms20092339 (2019).

24 Curtis, M. D. & Grossniklaus, U. A gateway cloning vector set for high-throughput functional analysis of genes in planta. Plant Physiology 133, 462–469, doi:DOI 10.1104/pp.103.027979 (2003).

25 Decaestecker, W. et al. CRISPR-TSKO: A Technique for Efficient Mutagenesis in Specific Cell Types, Tissues, or Organs in Arabidopsis. Plant Cell 31, 2868–2887, doi:10.1105/tpc.19.00454 (2019).

26 Schürholz, A. K. et al. A Comprehensive Toolkit for Inducible, Cell Type-Specific Gene Expression in Arabidopsis. Plant Physiology 178, 40–53, doi:10.1104/pp.18.00463 (2018).

27 Lampropoulos, A. et al. GreenGate - A Novel, Versatile, and Efficient Cloning System for Plant Transgenesis. Plos One 8, doi:ARTN e8304310.1371/journal.pone.0083043 (2013).

28 Waadt, R., Krebs, M., Kudla, J. & Schumacher, K. Multiparameter imaging of calcium and abscisic acid and high-resolution quantitative calcium measurements using R-GECO1-mTurquoise in Arabidopsis. New Phytologist 216, 303–320, doi:10.1111/nph.14706 (2017).

29 Dragwidge, J. M. & Van Damme, D. Protein phase separation in plant membrane biology: more than just a compartmentalization strategy. Plant Cell 35, 3162–3172, doi:10.1093/plcell/koad177 (2023).

30 Lakatos, L., Szittya, G., Silhavy, D. & Burgyán, J. Molecular mechanism of RNA silencing suppression mediated by p19 protein of tombusviruses. Embo J 23, 876–884, doi:10.1038/sj.emboj.7600096 (2004).

31 Li, X. Y. et al. A distinct endosomal Ca/Mn pump affects root growth through the secretory process. Plant Physiology 147, 1675–1689, doi:10.1104/pp.108.119909 (2008).

32 Van Leene, J. et al. A tandem affinity purification-based technology platform to study the cell cycle interactome in. Molecular & Cellular Proteomics 6, 1226–1238, doi:10.1074/mcp.M700078-MCP200 (2007).

33 Bechtold, N. & Pelletier, G. In planta Agrobacterium-mediated transformation of adult Arabidopsis thaliana plants by vacuum infiltration. Methods Mol Biol 82, 259–266, doi:10.1385/0-89603-391-0:259 (1998).

34 Dejonghe, W. et al. Disruption of endocytosis through chemical inhibition of clathrin heavy chain function. Nat Chem Biol 15, 641–+, doi:10.1038/s41589-019-0262-1 (2019).

35 Grones, P. et al. The endocytic TPLATE complex internalizes ubiquitinated plasma membrane cargo (vol 8, pg 1467, 2022). Nature Plants 9, 191–191, doi:10.1038/s41477-023-01344-w (2023).

36 Meier, F. et al. Online Parallel Accumulation-Serial Fragmentation (PASEF) with a Novel Trapped Ion Mobility Mass Spectrometer. Mol Cell Proteomics 17, 2534–2545, doi:10.1074/mcp.TIR118.000900 (2018).

37 Tyanova, S., Temu, T. & Cox, J. The MaxQuant computational platform for mass spectrometry-based shotgun proteomics. Nature Protocols 11, 2301–2319, doi:10.1038/nprot.2016.136 (2016).

38 Kourelis, J. et al. A homology-guided, genome-based proteome for improved proteomics in the alloploid. Bmc Genomics 20, doi:ARTN 72210.1186/s12864-019-6058-6 (2019).

39 Updegraff, D. M. Semimicro determination of cellulose in biological materials. Anal Biochem 32, 420–424, doi:10.1016/s0003-2697(69)80009-6 (1969).

40 Kumar, M. & Turner, S. Protocol: a medium-throughput method for determination of cellulose content from single stem pieces of Arabidopsis thaliana. Plant Methods 11, 46, doi:10.1186/s13007-015-0090-6 (2015).

41 Grefen, C., Lalonde, S. & Obrdlik, P. Split-ubiquitin system for identifying proteinprotein interactions in membrane and full-length proteins. Curr Protoc Neurosci Chapter 5, Unit 5 27, doi:10.1002/0471142301.ns0527s41 (2007).

