## Supplementary Figures for "Implementation of pupylation-based proximity labelling in plant biology reveals regulatory factors in cellulose biosynthesis"

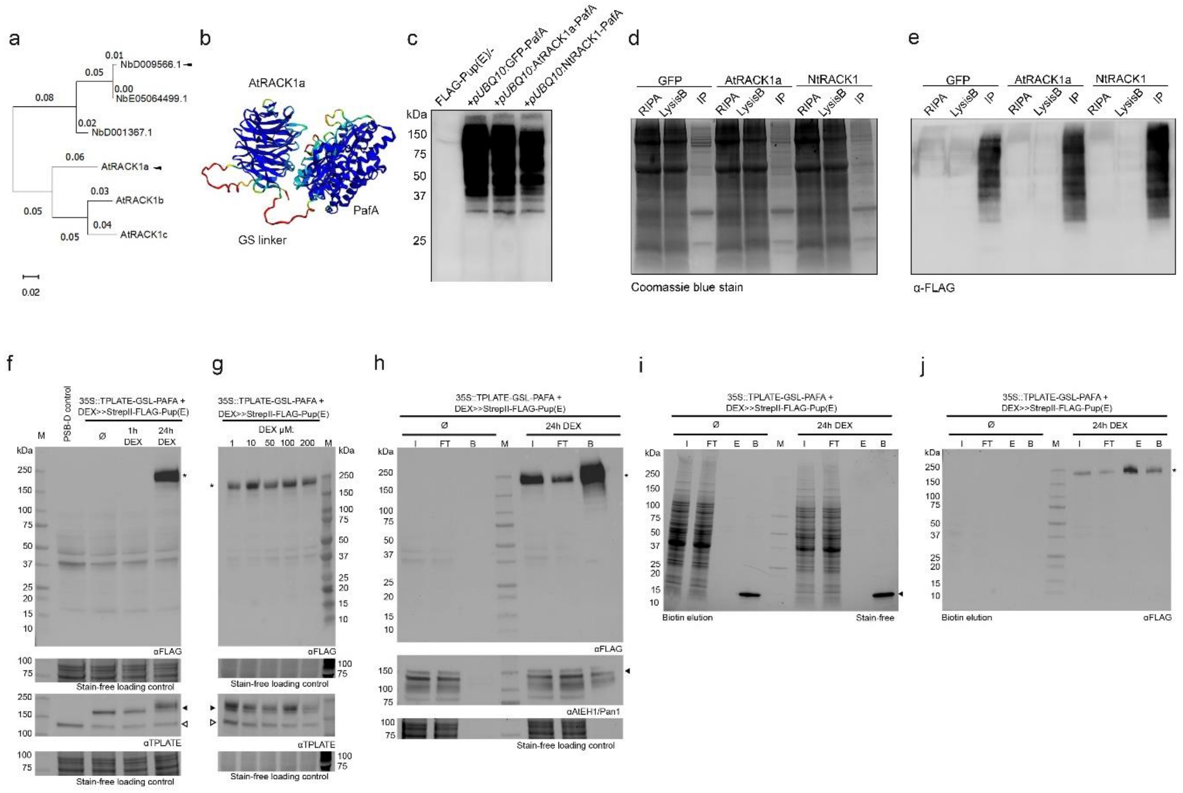


### Fig. S1: RACK1 and TPLATE pupylation.

**a**, A phylogenetic tree comparing the RACK proteins from tobacco and Arabidopsis. **b**, Glycine-Serine Linkers (GGGGSGGG) was introduced between the PafA and AtRACK1a with structure predicted by Alphafold2 (https://alphafold.ebi.ac.uk/). **c**, Western blot to detect the FLAG-Pup(E) labelled proteins from RACK1 transient expression in tobacco leaves. **d, e**, Pulldown of FLAG-Pup(E) labelled proteins. Coomassie blue stain indicated the total proteins in lanes (c) and the same amount of sample was loaded to detect the FLAG-Pup(E) labelled proteins with FLAG antibody (d). **f**, Inducible expression of StrepII-FLAG-Pup(E) after one hour and after 24 hours of 50µM dexamethasone (DEX) treatment in PSB-D cells constitutively expressing TPLATE-GSL-PafA, visualized using WB detection of Pup(E) via anti-FLAG and TPLATE via anti-TPLATE. The open arrowhead points to the endogenous TPLATE, while the closed arrowhead indicates the tagged form with PafA. Note that the band of the tagged form becomes blurry after 24h of induction, suggestive of post-translational modification with Pup(E). **g**, expression analysis of StrepII-FLAG-Pup(E) in the presence of TPLATE-GSL-PafA following a 24-hour treatment with different concentrations of dexamethasone (DEX). The 10µM concentration yielded the strongest pupylation level and was retained for further experiments. The open and closed arrowheads indicate the endogenous TPLATE as well as the tagged form. **h**, Pulldown experiment of StrepII-tagged proteins in PSB-D cells expressing TPLATE-GSL-PafA following a mock or 50µM dexamethasone treatment to induce expression of StrepII-FLAG-Pup(E). Next to the anti-FLAG detection, we also detected AtEH1/Pan1 as a proxy for complex incorporation of the PafA-tagged bait. **i j**, Pulldown experiment of StrepII-tagged proteins in PSB-D cells expressing TPLATE-GSL-PafA following a mock or a 10µM dexamethasone treatment (DEX) to induce expression of StrepII-FLAG-Pup(E), StrepII-tagged proteins (asterisk) were eluted from the beads using 50mM biotin. Strep-TactinXT® (black arrowhead) remains in the bead fraction and does not visibly contaminate the eluate.

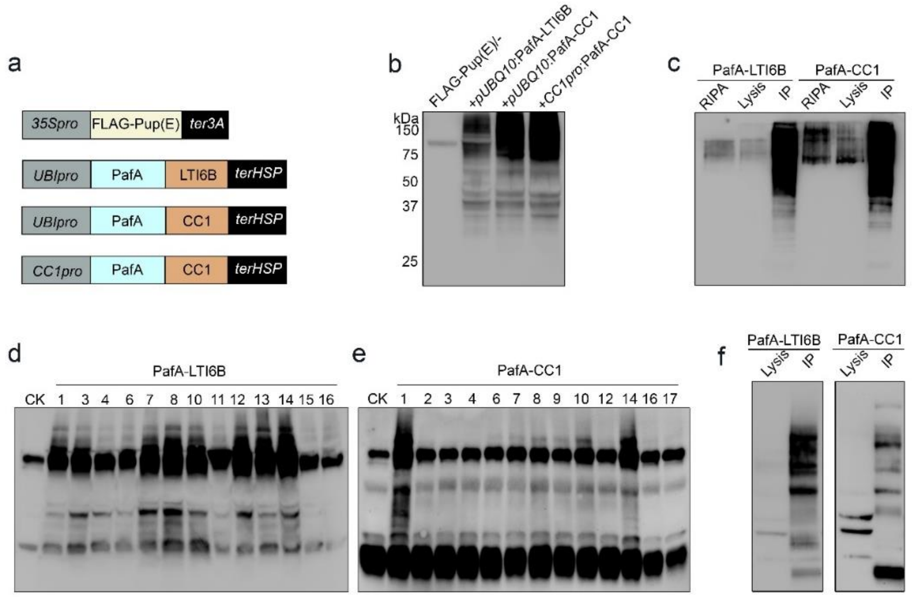


**Fig. S2: CC1 enrichment via pupylation.**

**a**, Vector design to express the FLAG-Pup(E), and PafA fused LTI6B/CC1.**b**, Western blot to detect the FLAG-Pup(E) labelled proteins from transient expression in tobacco leaves. Western blot was probed with anti-FLAG antibody. **c**, Pulldown of FLAG-Pup(E) labelled proteins from tobacco leaves and detection of the proteins using FLAG antibody. **d e**, Western blot to screen transgenic Arabidopsis that expressed the PUP-IT system with LTI6B (d) and CC1 (e). **f**, Pulldown of FLAG-Pup(E) labelled proteins in Arabidopsis, and subsequent detection of pupylated proteins using a FLAG antibody.

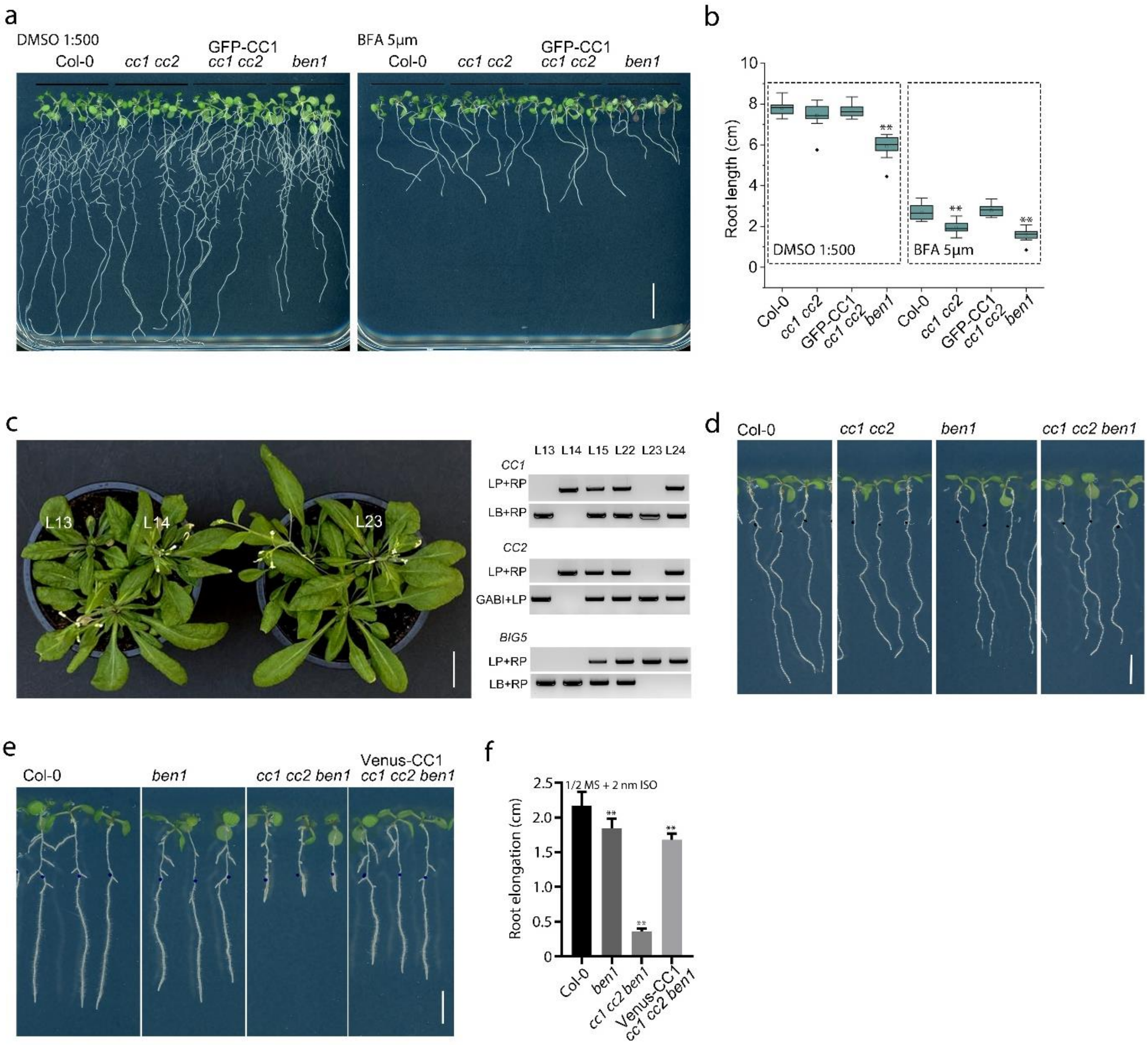


**Fig. S3: BEN1 affects trans-Golgi function and cellulose synthesis.**

**a,** Col-0 and CC1, BEN1 transgenic lines grown on media with DMSO (control) or BFA for seven days. Scale bar = 1 cm (a). **b,** Root length measurements from seedlings as those in a (Student t-test, $n > 10$, $**P < 0.01$). **c**, Genotyping to identify the *cc1 cc2 ben1* triple mutant, scale bar = 2 cm. **d, e,** Col-0 and CC1, BEN1 transgenic lines grown on DMSO (control) or isoxaben supplemented media. Scale bar = 0.5 cm. **f,** Root length measurements from seedlings as those in e (Student t-test, $n > 10$, $**P < 0.01$).
